# Structural elements that modulate the substrate specificity of plant purple acid phosphatases: avenues for improved phosphorus acquisition in crops

**DOI:** 10.1101/749564

**Authors:** Daniel Feder, Ross P. McGeary, Natasa Mitić, Thierry Lonhienne, Agnelo Furtado, Benjamin L. Schulz, Robert J. Henry, Susanne Schmidt, Luke W. Guddat, Gerhard Schenk

## Abstract

Phosphate acquisition by plants is an essential process that is directly implicated in the optimization of crop yields. Purple acid phosphatases (PAPs) are ubiquitous metalloenzymes, which catalyze the hydrolysis of a wide range of phosphate esters and anhydrides. While some plant PAPs display a preference for ATP as the substrate, others are efficient in hydrolyzing phytate or 2-phosphoenolpyruvate (PEP). PAP from red kidney bean (rkbPAP) is an efficient ATP- and ADPase, but has no activity towards phytate. The crystal structure of this enzyme in complex with an ATP analogue (to 2.20 Å resolution) provides insight into the amino acid residues that play an essential role in binding this substrate. Homology modelling was used to generate three-dimensional structures for the active sites of PAPs from tobacco (NtPAP) and *Arabidopsis thaliana* (AtPAP12 and AtPAP26) that are efficient in hydrolyzing phytate and PEP as substrates, respectively. In combination with substrate docking simulations and a phylogenetic analysis of 49 plant PAP sequences (including the first PAP sequences reported from *Eucalyptus*), several active site residues were identified that are important in defining the substrate specificities of plant PAPs. These results may inform bioengineering studies aimed at identifying and incorporating suitable plant PAP genes into crops to improve phosphorus use efficiency. Organic phosphorus sources increasingly supplement or replace inorganic fertilizer, and efficient phosphorus use of crops will lower the environmental footprint of agriculture while enhancing food production.

## Introduction

Phosphorus is a fundamental nutrient for living organisms. While animals rely on their diet for an adequate uptake of this element, plants have developed acquisition mechanisms that facilitate the absorption of phosphorus, mainly in the form of inorganic phosphate, from the rhizosphere. The concentration of inorganic ortho-phosphate in the soil is low (0.1–10 μM) [1], but organic substrates such as inositol phosphate, phospholipids, nucleotides, organic polyphosphate, phytate ((1*R*,2*r*,3*S*,4*S*,5*s*,6*S*)-cyclohexane-1,2,3,4,5,6-hexaylhexakis[dihydrogen(phosphate)]), and PEP (2-phosphoenolpyruvate) offer alternative sources of phosphorus and account for 30-90% of the soil phosphorus pool [2–4]. Phytate serves a range of biological functions in plants, most notably as a phosphorus and energy store [5]. PEP is an essential precursor in plants for the synthesis of chorismate in the shikimate pathway [6]. Consequently, plants have evolved enzymes such as phytases that hydrolyze phosphate ester bonds in phytate [1,7,8], and PEP phosphatases (PEPases) that cleave the phosphate group from PEP [9]. Another group of enzymes associated with phosphate acquisition by plants are the purple acid phosphatases (PAPs), metal ion-dependent hydrolases that are capable of liberating phosphate from a wide range of phosphorylated substrates [10]. The upregulation of intracellular and secreted PAPs is a universal plant response to phosphate deprivation, and transgenic PAP expression is considered as an avenue for improved phosphorus acquisition by crops (Tran, Hurley, *et al.*, 2010; Tran, Qian, *et al.*, 2010). More generally, improving the access of crops to organic phosphorus is desirable as re-purposed organic wastes, including manures, compost and agricultural residues, are increasingly used to supplement or replace inorganic (mineral) fertilizers. Identifying effective root-derived phosphatases and boosting their presence in the rhizosphere may be exploited in agriculture as these enzymes may lessen the demand for fertilizers, thus enhancing sustainability and increasing efficiency. Globally, the phosphorus fertilizer utilization efficiency is estimated to be 20-50% and may be as low as 16% for cereals, which account for 61% of the total harvested cropland [13]. Thus, phosphorus-associated environmental footprints come at a great cost to water sources, farmers and the ecosystem in particular through eutrophication [14] and/or pollution of the hydrosphere [15].

PAPs have been characterized from a number of sources, including plant species, *e.g. Arabidopsis thaliana* [12, 16], red kidney bean [17, 18], soy bean [19], tobacco [20] and tomato [16, 21], as well as animal species, *e.g.* bovine [22], human [23, 24] and pig [25, 26]. The majority of known plant PAPs form homodimers of ∼55 kDa subunits, while mammalian PAPs are ∼35 kDa monomers [10]. It is interesting to note, however, that small “mammalian-like” PAPs have been discovered in some plants [27, 28] and a “plant-like” PAP gene has been identified in the human genome [29]. PAPs require metal ions for their function, and despite diversity with respect to their overall sequences, all PAPs characterized so far use the same seven amino acid residues in their active sites to coordinate the two catalytically essential metal ions (**Fig. 1**). PAPs are the only known hydrolases that require a heterovalent bimetallic center of the form Fe(III)-M(II) (where M = Fe, Zn or Mn) to promote catalytic activity (although an active di-Mn(II) form could be generated *in vitro*) [10,19,30–32]. The Fe(III) and its tyrosine ligand form a charge transfer complex absorbing at ∼560 nm, thus leading to the characteristic purple color of the enzyme [10].

**Figure 1:**
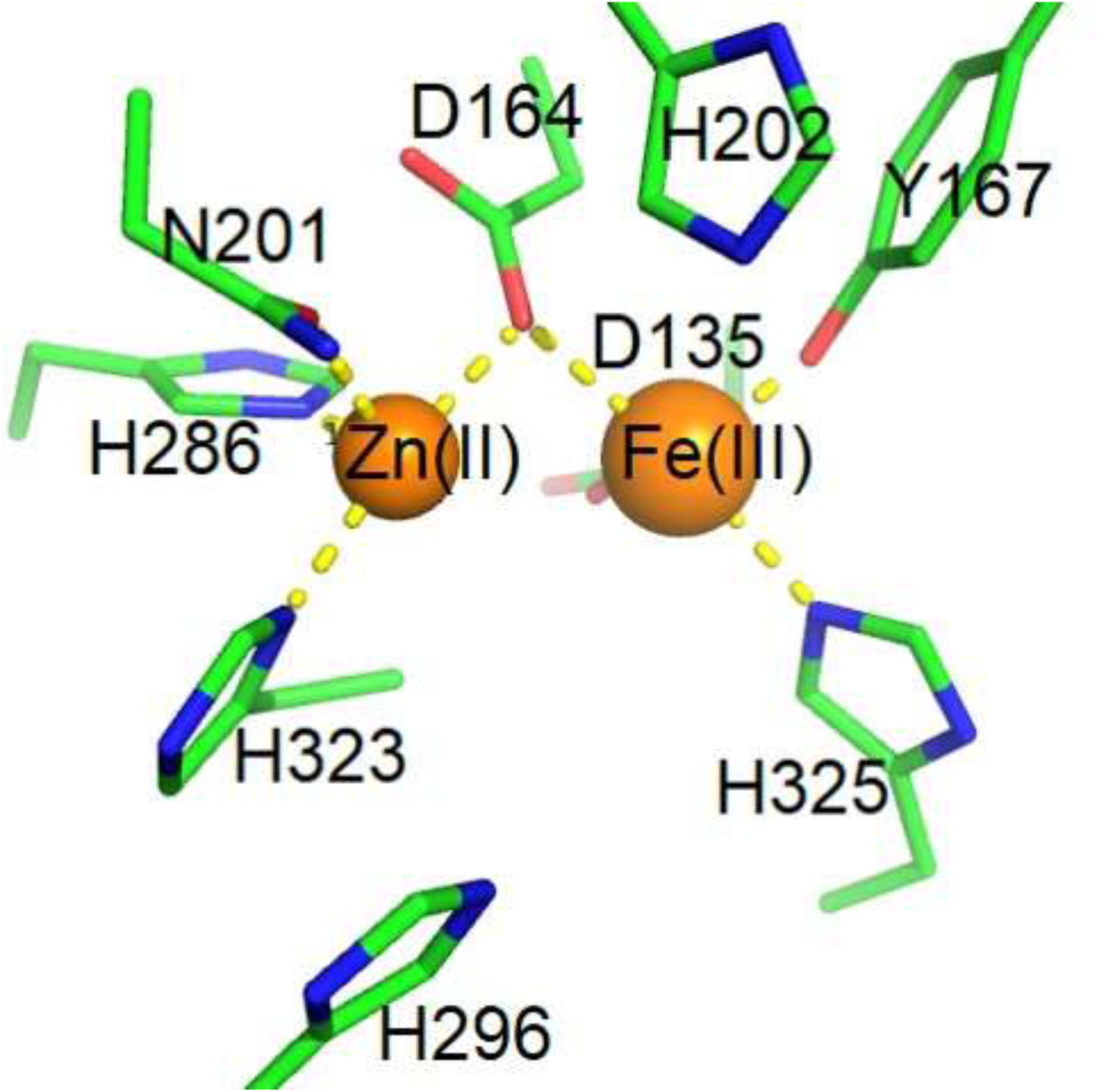
Illustration of the PAP active site (green carbons; numbering according to the sequence of rkbPAP). The interaction between Y167 and Fe(III) leads to the characteristic purple color of PAP [10, 30]. Bonds between amino acid ligands and metal ions are indicated as dashed yellow lines.

PAPs are characterized by their capability to hydrolyze a broad range of substrates including adenosine 5’-triphosphate (ATP), adenosine 5’-diphosphate (ADP), phosphotyrosine and pyrophosphate [10]. The synthetic substrate *para*-nitrophenyl phosphate (*p-*NPP) is frequently used for mechanistic studies [10,30,33]. In animals, the ability of PAP to dephosphorylate proteins such as osteopontin and bone sialoprotein may align with the enzyme’s proposed function in bone metabolism [34]. It could be demonstrated that proteolytically-activated human PAP is a potent ATPase, raising the possibility that this enzyme may also play an important role in energy metabolism [24]. Furthermore, animal PAPs are bifunctional, at least *in vitro*, since they can also act as peroxidases [35–37].

In plants, the assignment of precise role(s) for PAPs is obscured due to the occurrence of multiple isoforms of both the smaller and larger variants of this enzyme. In *A. thaliana*, 29 PAPs were identified by genome annotation [27], and the emerging picture indicates that different PAP isoforms may have different functions, a least in this plant. For instance, isoform 26 (AtPAP26) is strongly upregulated on root surfaces and in cell walls of phosphorus-starved seedlings, and its function cannot be compensated by other AtPAP isoforms [12,38–40]. The ortholog of AtPAP26 in rice (OsPAP26) is also upregulated during phosphate deprivation and thus may fulfil a similar crucial role in phosphorus metabolism [41]. Importantly, in the context of the role of plant PAPs in P acquisition, at least some representatives (*e.g.* from soybean and tobacco) can hydrolyze phytate [8, 42]. The best studied plant PAP to date is from red kidney bean (rkbPAP) [17]. Although a considerable amount of structural and mechanistic data is available, the precise physiological function(s) of this enzyme is still subject to debate [43–45]. Motivated by this gap in knowledge we identified specific structural elements that play an important role in determining the selectivity of plant PAPs for substrates. The insight gained will inform future bioengineering studies that will use PAPs to enhance nutrient uptake by plants, an essential step towards improving sustainable food production for a growing global population.

## Methods and Methods

### Enzyme preparation and purification

rkbPAP was purified following a previously published protocol [17]. Briefly, red kidney beans (*Phaseolus vulgaris*) were ground in a Waring blender and suspended in 0.5 M sodium chloride. The suspension was filtered through a muslin cloth, followed by ethanol fractionation and ammonium sulfate precipitation. Purification was carried out by ion-exchange chromatography using a CM-cellulose column, followed by gel filtration on a Sephadex S-300 column. The resulting preparation was concentrated to 23.8 mg/mL using a Millipore Amicon centrifugal concentrator and stored at 4°C in 0.5 M sodium chloride.

### Determination of kinetic parameters

For standard assays of the catalytic properties of rkbPAP, *p*-NPP was used as the substrate. Product formation at pH 4.9 was measured at λ = 390 nm and ε = 342.9 M^−1^cm^−1^ [46]. However, in order to provide a tool to assess the catalytic properties of plant PAPs for substrates lacking suitable chromophores (*e.g.* ATP, ADP and phytate) an assay was established that employs isothermal titration calorimetry (ITC) similar to a protocol that was recently developed for a group of related metallohydrolases [47]. Reactions were initiated by injection of 5 – 10 µL rkbPAP (0.16 µM stock solution) into mixtures with different concentrations of substrates (see Supplementary Section for further details). The reaction with *p*-NPP was used to calibrate the methodology. k_cat_ (s^−1^) and *K*_m_ (mM) were estimated using the Michaelis-Menten equation (where V_max_ = k_cat_ × [E]_0_).

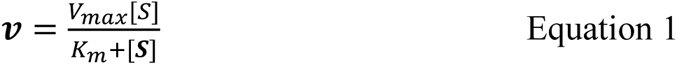

### Multiple sequence alignment and phylogenetic analysis

Multiple sequence alignments and phylogenetic analysis were undertaken using Clustal-O [48] and displayed with ESPript 3.0 [49] using PAP and predicted/hypothetical PAP sequences from 49 different plants (**Fig. S1**). Plant PAP sequences were obtained by performing a BLAST search with the rkbPAP sequence as the query and limiting the search to plant species that have applications in agriculture or biotechnology. The protein sequences from the BLAST search were identified as PAPs based on their characteristic seven metal-ligating residues and the two histidine residues known to stabilize the phosphate group of the substrate (His296 And His202 in rkbPAP). “Mammalian-like” plant PAPs were excluded from the analysis. The phylogenetic tree was generated using iTOL [50].

### Crystallization and cryprotection

Crystals of rkbPAP were grown by the hanging drop diffusion method using a previously established crystallization medium [45]. Once formed the crystals were transferred into a drop containing cryoprotectant solution (containing 0.1 M sodium citrate pH 5.0, 0.1 M lithium chloride, 25% polyethylene glycol 3,350, 20% isopropyl alcohol, 10% glycerol, 5 mM vanadate and 5 mM ADP) and allowed to soak for four days before cryo cooling in liquid nitrogen.

### Data collection and analysis

Data were collected at the Australian Synchrotron MX1 beam-line and were scaled and merged using XDS [51]. Structures were solved by difference Fourier techniques using a previously reported structure of rkbPAP (PDB code: 4DHL [52]) as the starting model. Final refinements were undertaken using PHENIX [53]. WinCoot [54] was used for electron density fitting and model building. All surface figures, as well as the stick models of structures were generated using Molegro Virtual Docker (MVD) [55]. Omit map figures were created with CCP4MG [56] and stick figures with PyMOL [57] unless stated otherwise.

### In-silico docking and homology modeling

Docking studies were undertaken with MVD using the MolDock SE algorithm. The coordinates of the crystal structure of rkbPAP-ADP-metavandate obtained in this study (PDB: 6PY9) were used for docking of ADP and ATP to rkbPAP. ADP-metavanadate was omitted from the active site for the docking studies. The ligand search space was confined to a sphere of 9 Å radius originating from the metal center of rkbPAP. The chromophoric Fe(III) in the active site was assigned the charge of +3. Prior to setting up docking simulations, water molecules were removed from the coordinate file using MVD. The poses with the most negative MolDock, Rerank and HBond scores with no steric/electrostatic clashes with the protein were selected as they represent the most preferred and stable energetic states of the substrate bound to the protein. Models of a PAP from tobacco (NtPAP) and one isoform from *A. thaliana* (AtPAP12) were generated *in silico* with WinCoot [54] by mutating relevant residues in the structure of rkbPAP with those corresponding to the NtPAP or AtPAP12 sequences and fitting them to the structure with a rotamer conformation consistent with that for the crystal structure of rkbPAP. In order to create a model of the *A. thaliana* PAP isoform AtPAP26 the structure of sweet potato PAP (IbPAP2) [58] was used due to the high degree of similarity. For docking studies, the side chains of the residues that were mutated in rkbPAP (to their respective NtPAP, AtPAP12 or AtPAP26 counterparts) were allowed to be flexible during the docking simulation. All other settings were similar to those in the docking studies with rkbPAP and the substrates ADP and ATP. Superimpositions of structures and mutant models were carried out with MVD.

## Results and Discussion

The biological functions of PAPs are diverse and in plants the assignment of specific roles is complicated by the fact that these enzymes occur in multiple isoforms that may differ in their substrate preference [11,12,39,59]. Some PAPs act as phytases (so called purple acid phytases), including PAPs from *Nicotiana tabacum* (tobacco), *Oryza sativa* (rice) and *Glycine max* (soybean) [20,42,60]. Other PAPs prefer PEP as their substrate, including two PAP isoforms from *A. thaliana*, *i.e.* AtPAP12 and AtPAP26 [12], while others are known to be ATPases, including AtPAP10, a PAP from sweet potato (IbPAP1) and rkbPAP [17,43,61,62]. Despite being enzymatically the most-characterized plant PAP, the precise physiological role(s) of rkbPAP remains unknown.

A difficulty in comparing the efficiency of PAPs towards different substrates is the absence of useful chromophoric markers that facilitate rapid and continuous monitoring of the reaction progress with substrates such as ADP, ATP or phytate. While discontinuous assays that record the rate of formation of the reaction product phosphate are available, they are time-consuming and require considerable amounts of enzyme [63]. Recently, we established a rapid and continuous assay to measure the reaction rates of several hydrolases, including organophosphate-degrading hydrolases and the cyclic nucleotide disesterase Rv0805 [64–67] by monitoring the enthalpy change associated with the cleavage of the relevant ester bonds [47]. We have now established a similar assay for PAPs using the substrate *p*-NPP as calibrant to compare the catalytic parameters obtained from the calorimetric and spectroscopic assays (see Supplementary Section and **Fig. S2** for details). The calorimetrically determined catalytic parameters (*k_cat_* = 190 s^−1^ and *K*_m_ = 4 mM) are in good agreement with the corresponding values determined spectrophotometrically (180 s^−1^ and 2.6 mM, respectively). This assay can easily be adapted for other substrates of PAPs as well and will be a useful tool to establish the substrate preference of diverse PAPs.

### Crystallographic investigations

In a previous study we demonstrated that the soaking of rkbPAP crystals in a cryosolution containing both vanadate and adenosine leads to adenosine divanadate (ADV) being formed in a complex that mimics the transition state of ADP hydrolysis to AMP by PAP [18]. No monovanadate derivatives of adenosine (AMV) were observed. Here, we expanded this study by crystallizing rkbPAP in the presence of ADP and vanadate. Although ADP is both a substrate and product of the PAP-catalyzed reaction we hypothesized that the presence of the potent PAP inhibitor vanadate may lead to the *in crystallo* formation of ADP metavanadate (ADPV), which forms spontaneously in solution and has been used extensively as a model for ATP in ATP-binding proteins [68]. Relevant crystallographic parameters are summarized in **Table 1**. There are two dimers (subunits A-D) in each asymmetric unit of the crystal. The overall fold of the four polypeptide chains strongly resembles those of the previously determined structures of free rkbPAP [69] and the phosphate-sulfate- and vanadate-bound complexes [18,44,46] of this enzyme (rmsd values for all Cα atoms <0.353 Å). For initial fitting of the ligands in the active site, Polder omit maps were chosen due to their ability to provide enhanced electron density when compared to simulated annealing omit maps [70]. Due to the best quality electron density data for subunit A we will focus only on this subunit to describe the structure of the rkbPAP-ligand complex that formed in the crystallization experiment.

**Table 1.**
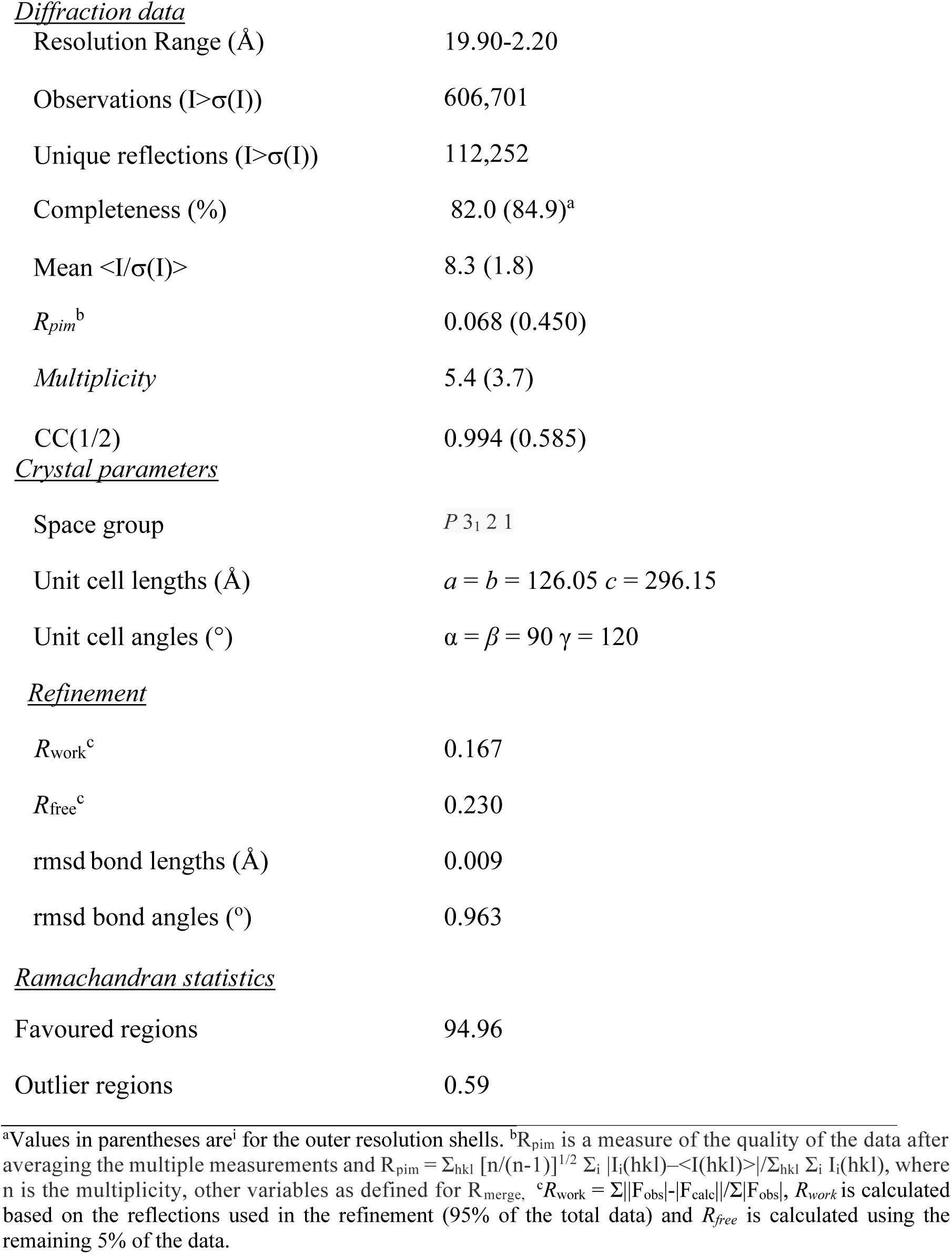
Data collection and refinement statistics for the rkbPAP-ADP-metavanadate complex (rkbPAP-ADPV).

Diffraction data
|  |  |
| --- | --- |
| Resolution Range (Å) | 19.90-2.20 |
| Observations ($I > \sigma(I)$) | 606,701 |
| Unique reflections ($I > \sigma(I)$) | 112,252 |
| Completeness (%) | 82.0 (84.9)<sup>a</sup> |
| Mean $\langle I/\sigma(I) \rangle$ | 8.3 (1.8) |
| $R_{\text{pim}}^b$ | 0.068 (0.450) |
| Multiplicity | 5.4 (3.7) |
| CC(1/2) | 0.994 (0.585) |
| Crystal parameters |  |
| Space group | $P 3_1 2 1$ |
| Unit cell lengths (Å) | $a = b = 126.05$ $c = 296.15$ |
| Unit cell angles (°) | $\alpha = \beta = 90$ $\gamma = 120$ |
| Refinement |  |
| $R_{\text{work}}^c$ | 0.167 |
| $R_{\text{free}}^c$ | 0.230 |
| rmsd bond lengths (Å) | 0.009 |
| rmsd bond angles (°) | 0.963 |
| Ramachandran statistics |  |
| Favoured regions | 94.96 |
| Outlier regions | 0.59 |

<sup>a</sup>Values in parentheses are<sup>i</sup> for the outer resolution shells. <sup>b</sup> $R_{\text{pim}}$ is a measure of the quality of the data after averaging the multiple measurements and $R_{\text{pim}} = \sum_{\text{hkl}} [n/(n-1)]^{1/2} \sum_i |I_i(\text{hkl}) - \langle I(\text{hkl}) \rangle| / \sum_{\text{hkl}} \sum_i I_i(\text{hkl})$ , where $n$ is the multiplicity, other variables as defined for $R_{\text{merge}}$ , <sup>c</sup> $R_{\text{work}} = \sum ||F_{\text{obs}}| - |F_{\text{calc}}|| / \sum |F_{\text{obs}}|$ , $R_{\text{work}}$ is calculated based on the reflections used in the refinement (95% of the total data) and $R_{\text{free}}$ is calculated using the remaining 5% of the data.

ADPV is bound to the binuclear metal center of rkbPAP (0.60 occupancy; average B factor of 55.56) (**Fig. 2a**)^1^. The oxygen atom O1 of the vanadate group (**Fig. 2b**) binds to Zn(II) (2.3 Å) and forms a hydrogen bond with the sidechain amino group of Asn201 (2.6 Å). O2 binds strongly to Fe(III) (2.0 Å) and O3 is in a metal-bridging position between Zn(II) (2.6 Å) and Fe(III) (2.5 Å). The phosphate of ADPV that is linked to the vanadate group (*i.e.* the proximal phosphate group) is stabilized through a hydrogen bond between O5 and His202 (3.1 Å) and a weaker hydrogen bond between O5 and Tyr365 (3.4 Å). O6 forms a hydrogen bond with His295 (3.0 Å). The second phosphate group (*i.e.* the distal phosphate group) forms a hydrogen bond between O8 and an adjacent water molecule (W1; 2.3 Å) that is H-bonded to a nitrogen in the guanidino group of Arg258 (2.4 Å). It is also noteworthy that the vanadate moiety adopts a four-coordinate tetrahedral geometry, in contrast to the five-coordinate trigonal bipyramidal conformation observed in the ADV complex of rkbPAP [18]. Thus, while the latter was proposed to mimic the transition state of catalysis, the conformation of the former mimics the substrate-bound form [45]. The ribose moiety does not appear to form significant hydrogen bonds or other strong interactions with the enzyme, with the closest contact to the oxygen in the ribose ring being a nitrogen from the guanidino group of Arg258 (3.8 Å). The ribose hydroxyls are not involved in any bonding interactions. This paucity of interactions explains why the ribose moiety is only partially observed in the Polder electron density map, consistent with the high B factors of C2 from the ribose ring and the oxygen atom in the 2’-ribose hydroxyl (B= 67.90 Å^2^ and 72.01 Å^2^, respectively), which are not covered by electron density. Water molecules are observed that form hydrogen bonds with nitrogen atoms in the adenine moiety (**Fig. 2b**); N2 is H-bonded to a water (W2, 3.0 Å) that is H-bonded to an oxygen in the sidechain carboxylate group of E299 (3.2 Å). N3 is H-bonded to a water (W3, 3.2 Å) that is H-bonded to the nitrogen in the backbone amine group of His296 (3.1 Å). N4 is H-bonded to a water (W4, 2.7 Å) that is H-bonded (3.2 Å) to the oxygen in the backbone carbonyl group of Asn294 (2.6 Å). N5 is stabilized by forming two hydrogen bonds, one with a water (W5, 3.2 Å) that is H-bonded to the backbone carbonyl of F297 (2.7 Å) and another with a water from the solvent (W6, 2.6 Å).

**Figure 2:**
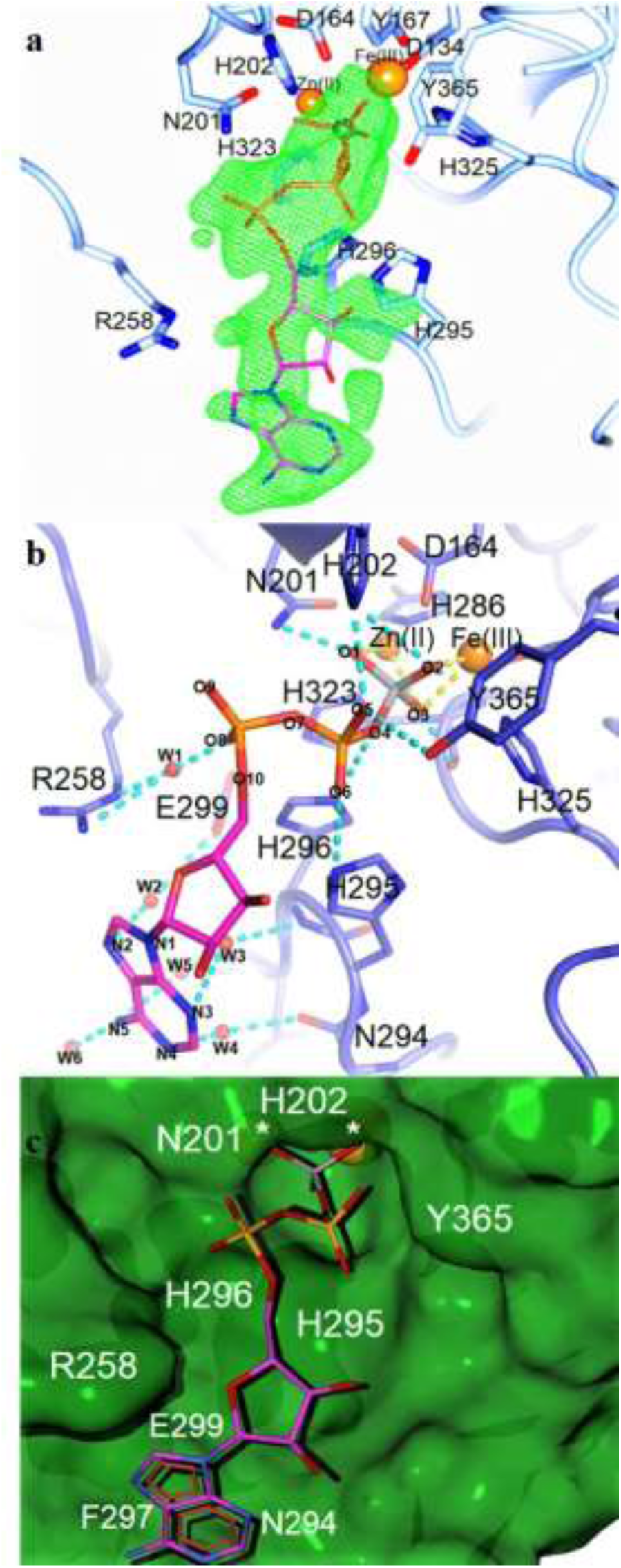
(a) Stick representation and Polder omit map (>3.1 σ) of ADPV (magenta carbons) in the active site of rkbPAP (light blue carbons). Fe(III) and Zn(II) are shown as orange spheres. The vanadium atom in the vanadate moiety of ADPV is shown as a gray sphere. (b) Cartoon and stick representation of ADPV (magenta carbons) in the active site of rkbPAP (light blue carbons). Fe(III) and Zn(II) are shown as orange spheres, V is in gray. Relevant oxygen and nitrogen atoms of the ligand are numbered (O1-O10 and N1-N5, respectively). Water molecules are shown as red spheres and labeled W1-W6. Hydrogen bonds are in dashed cyan and bonds to metal ions are in dashed yellow. (c) Surface and stick representation of the rkbPAP-ADPV complex. Metal ions are shown as orange spheres and marked with an asterisk.

### Sequence analysis, phylogenetic characterization and homology modeling of plant PAPs

It is important to point out that PAPs are capable of hydrolyzing a broad range of monoester substrates. However, while rkbPAP is a potent ATPase/ADPase and PEPase, this enzyme, in contrast to other plant PAPs, does not act as a phytase. In order to identify structural features in plant PAPs that play a crucial role in guiding substrate specificity a comprehensive multiple sequence alignment and a phylogenetic analysis were carried out using 49 sequences (Figs 3 **& S1**). Amongst those sequences are known phytases (*i.e.* NtPAP [20], OsPHY1 [71] and GmPHY [60]) and PEPases (AtPAP12 and AtPAP26 [12, 72]); similar to rkbPAP AtPAP10 [61] has been reported as an efficient ATPase but a poor phytase. It is also noted that while NtPAP acts as a phytase, it also accepts ATP and PEP as substrates [8].

**Figure 3:**
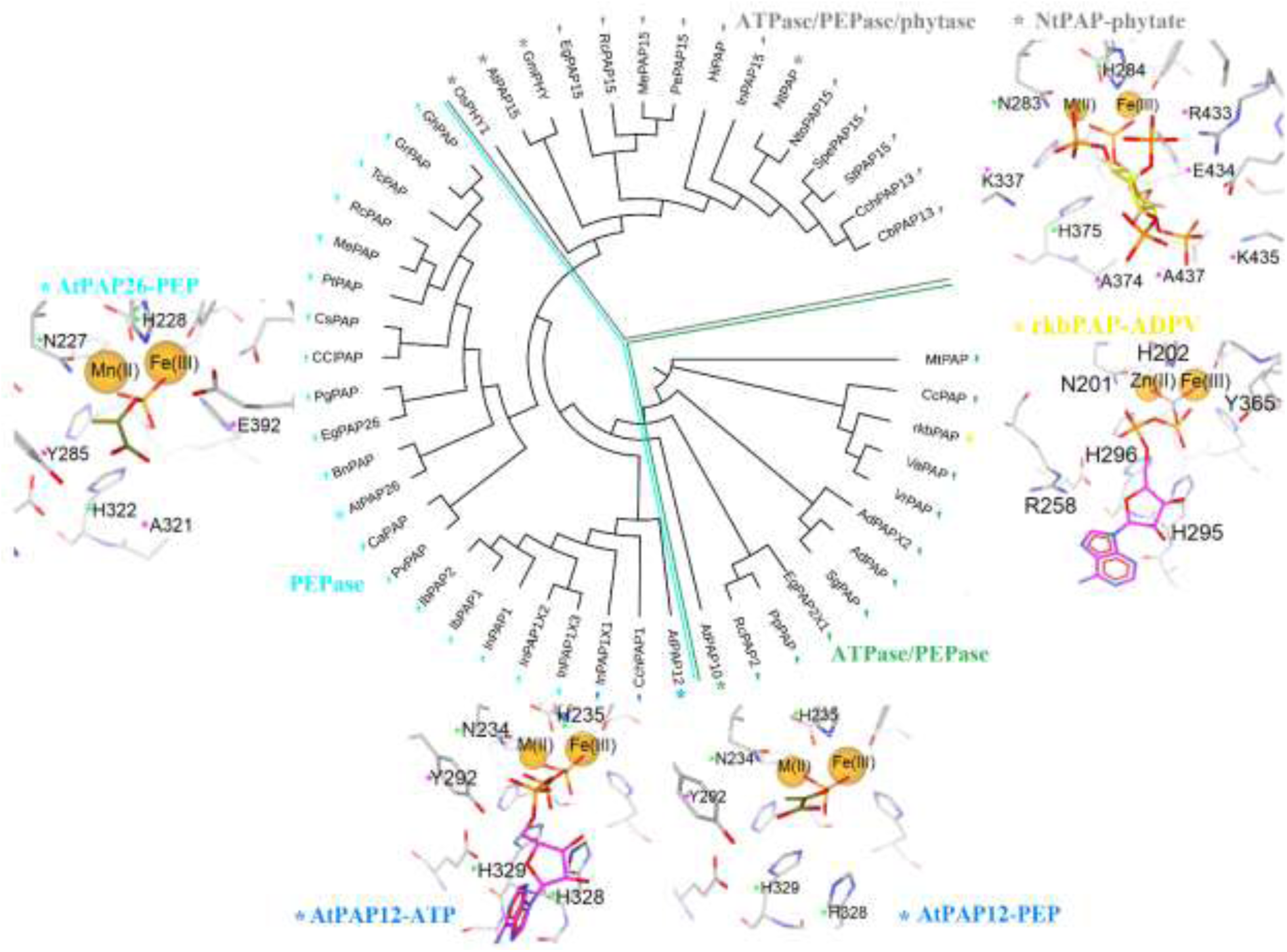
Phylogenetic tree analysis using 49 PAP sequences from agriculturally relevant plant species. PAPs with known substrate preference are marked with an asterisk (green for ATPase, cyan for PEPase, gray for ATPase/PEPase/phytase, blue for PEPase/ATPase and yellow for rkbPAP (ATPase/PEPase)). The sequences cluster according to their preferred substrates. The active sites of PAPs with varying substrate preference(s) are also shown (see text for details). Residues that are invariant are marked with a green asterisk and residues that differ relative to the rkbPAP sequence are marked with a magenta asterisk. Ad: *Arachis duranensis*; At: *Arabidopsis thaliana*; Bn: *Brassica napus*; Cb: *Capsicum baccatum*; Cc: *Cajanus cajan*; Cch: *Capsicum chinense*; Ccl: *Citrus clementina*; Ca: *Cicer arietinum*; Cs: *Citrus sinensis*; Eg: *Eucalyptus grandis*; Gh: *Gossypium hirsutum*; Gm: *Glycine max*; Gr: *Gossypium raimondii*; Hi: *Handroanthus impetiginosus*; Ib: *Ipomoea batatas*; In: *Ipomoea nil*; Me: *Manihot esculenta*; Mt: *Medicago truncatula*; Nt: *Nicotiana tabacum*; Nto: *Nicotiana tomentosiformis*; Os: *Oryza sativa*; Pe: *Populus euphratica*; Pg: *Punica granatum*; Pp: *Prunus persica*; Pt: *Populus trichocarpa*; Pv: *Phaseolus vulgaris* (French variant); Rc: *Ricinus communis*; Sg: *Stylosanthes guianensis*; Sp: *Solanum pennellii*; St: *Solanum tuberosum*; Tc: *Theobroma cacao*; Va: *Vigna angularis*; Vr: *Vigna radiate*.

The sequence alignment (Fig. S1) reveals that the seven metal-coordinating residues as well as the two histidine residues in the outer coordination sphere that play an important role in the catalytic mechanism of PAPs and/or binding interactions with the phosphate group of the substrate (His202 and His296 based on rkbPAP numbering) are invariant. However, plant PAPs with known substrate preference appear to cluster in distinct groups in the phylogenetic tree (**Fig. 3**). In order to correlate sequence variations with substrate preference homology models of NtPAP (a PAP that hydrolyzes ATP, PEP and phytate), AtPAP12 (a good PEP- and ATPase), and AtPAP26 (a PEPase with poor ATPase activity) were generated based on the crystal structures of rkbPAP (NtPAP and AtPAP12) or sweet potato PAP (AtPAP26). A comparison of the active site structures of the models is shown in **Fig. 4**. Subsequently, several representatives of biologically significant and/or agriculturally relevant substrates were docked into the respective active site models. The accuracy of this *in silico* approach was assessed by docking ADP and ATP into the active site of rkbPAP and comparing the outcome with the crystal structures of the rkbPAP-ADV [18] and rkbPAP-ADPV (**Fig. S3**) complexes. Irrespective of the identity of the docked substrate molecule (*i.e.* ADP or ATP) amino acid side chains of rkbPAP that participate in interactions with the substrates are Asn201, His202, Arg258, His295, His296 and Tyr365. These residues form a series of hydrogen bonds and electrostatic interactions with the phosphate groups of the substrates. For the modeled ADP complex the ribose group of the substrate is predicted to lack any strong interactions with rkbPAP, while the adenine ring is only weakly stabilized through a single hydrogen bond between a nitrogen atom (N3) and His295. This largely agrees with the crystal structure of the rkbPAP-ADV complex, where the absence of electron density for the adenine moiety in the adenosine group was explained by the weak interactions holding this moiety in place [18]. For the modelled ATP complex the ribose moiety is also predicted to lack significant interactions with the enzyme, but the adenine moiety is predicted to form a hydrogen bond between an adenine nitrogen (N5) and the backbone carbonyl oxygen of Phe297. Thus, despite the fact that docking was performed without the inclusion of water molecules, similar interactions were observed for ATP (**Fig. 5**) as in the structure of the rkbPAP-ADPV complex (**Fig. 2**). The combined docking data for the two rkbPAP complexes thus demonstrate that the *in silico* algorithm is successful in predicting the correct contact points for a substrate in the active site of a plant PAP. The docking scores (MolDock/Rerank scores [55], see **Table 2**) for ADP and ATP are -223.3/-150.0 and -337.7/-244.9 kcal/mol, respectively, indicating that the binding of these two substrates to the active site of rkbPAP is energetically highly favorable. Furthermore, the difference in the docking scores is mostly consistent with the observed affinities (based on Km values) for the different substrates tested, which indicate that rkbPAP prefers ATP over ADP. Subsequently, PEP and phytate were also docked into the active site of rkbPAP. The corresponding docking scores are -184.9/-122.4 and - 192.6/26, respectively. Thus, PEP is predicted to have a size-specific binding energy higher than that of ATP (ligand efficiencies (LEs) for the two substrates are -18.5 kcal/mol and - 10.9 kcal/mol per heavy atom, respectively). The most energetically favorable pose for phytate has clashes with most residues in the active site (**Fig. 5**), as reflected by its positive Rerank score, and consistent with the absence of detectable activity of rkbPAP towards this substrate.

**Figure 4:**
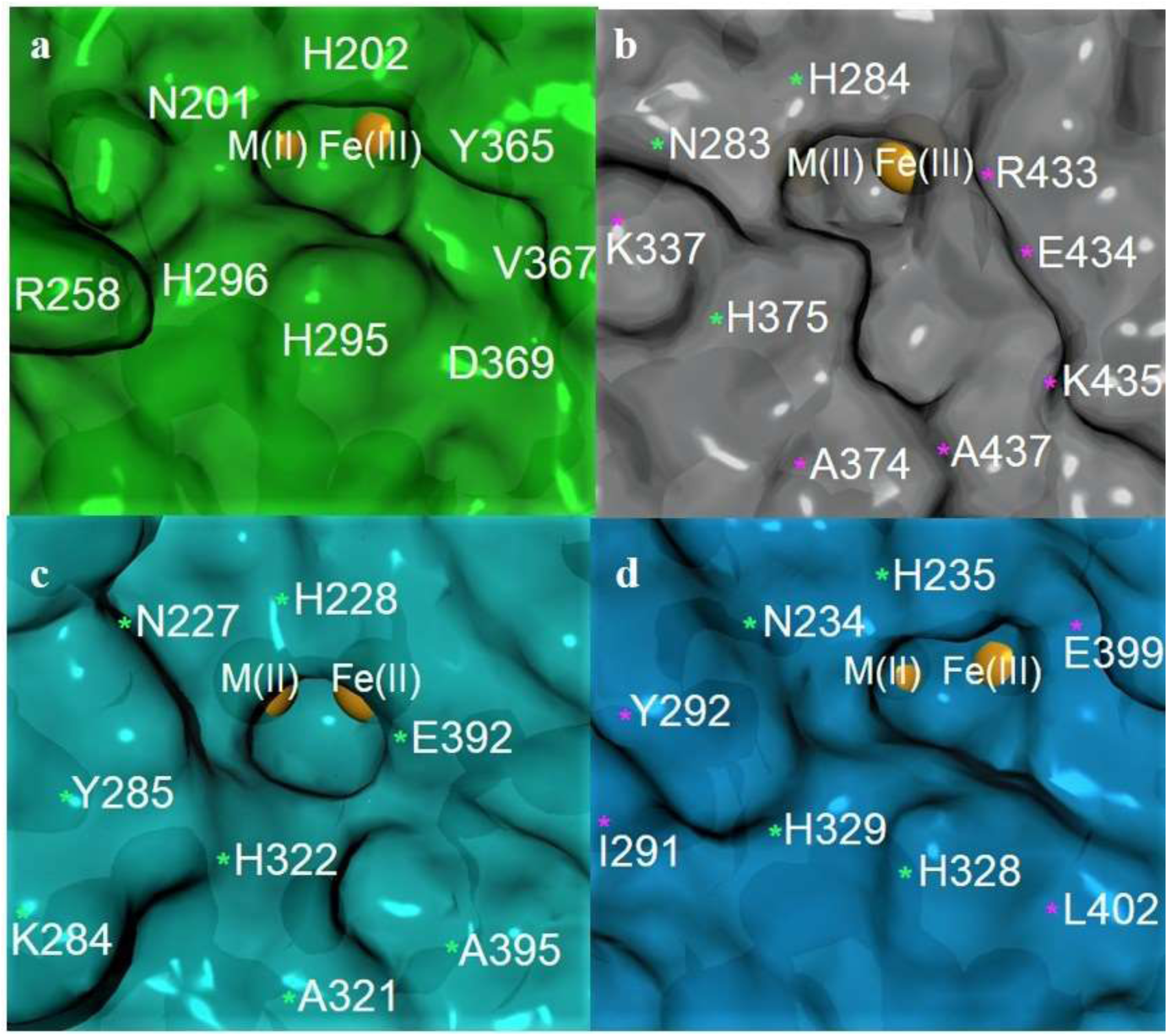
A comparison of the surfaces of the active sites of (a) rkbPAP (based on its crystal structure [52]), (b) NtPAP, (c) AtPAP26 and (d) AtPAP12. For the latter three enzymes their active sites were modelled as described in the text. In NtPAP, an elongated, narrow groove is present above the catalytic metal center that appears to be suitable for larger substrates such as phytate, while for AtPAP26 a small circular tunnel is formed above the metal center, suitable for smaller substrates such as PEP. In AtPAP12, the active site features structural elements from both PAP ATPases such as AtPAP10 and PEPases such as AtPAP26. Residues in NtPAP, AtPAP26 and AtPAP12 that differ from the corresponding residues in the original model are marked with a magenta asterisk; conserved residues relative to the original model are marked with a green asterisk.

**Figures 5:**
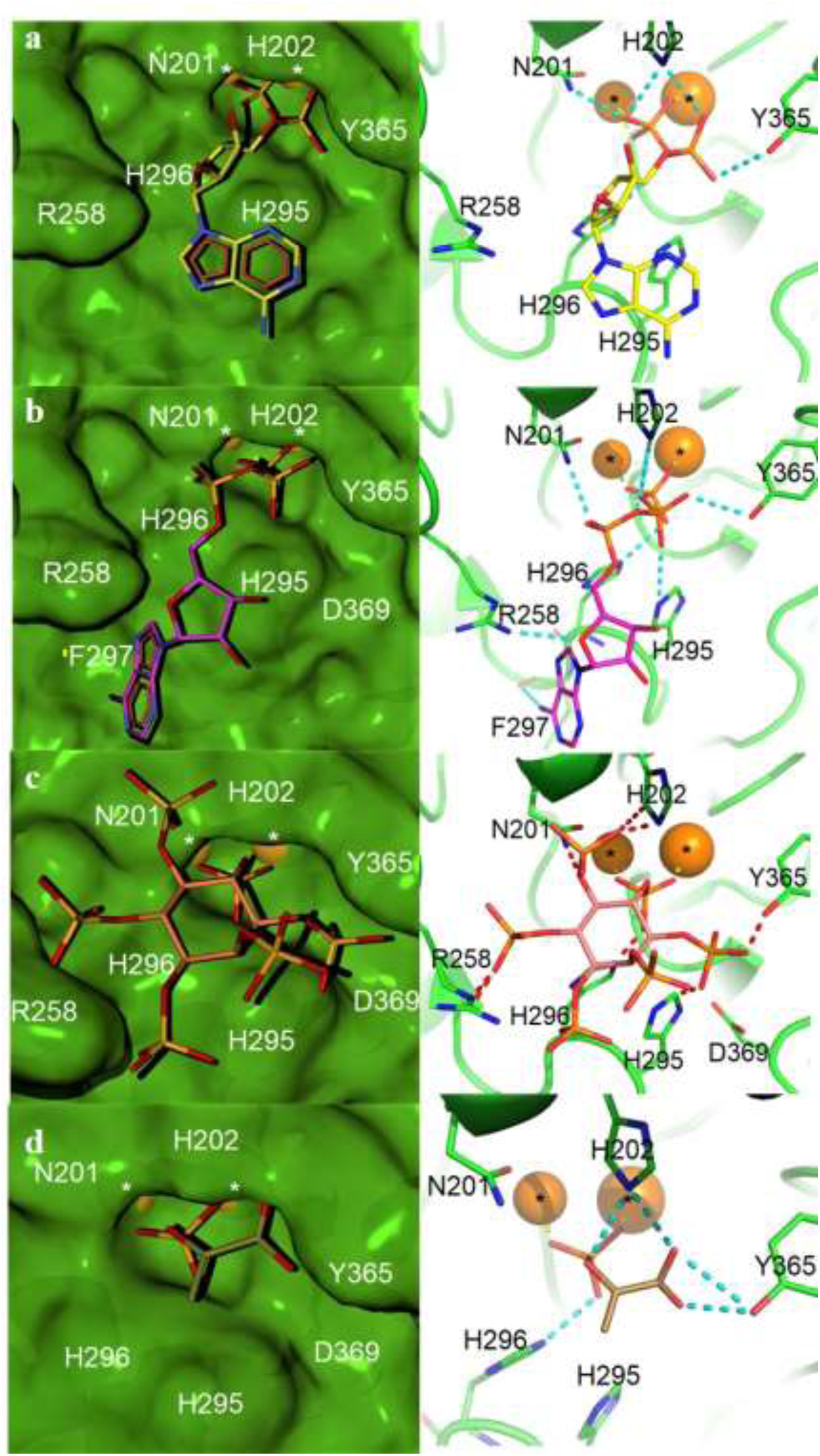
(left panel) Surface and (right panel) stick representations of the docking predictions of the binding modes of ADP (yellow carbons), ATP (magenta carbons), phytate (salmon carbons) and PEP (brown carbons) to rkbPAP (blue surface or carbons). Hydrogen bonds are indicated in dashed cyan and bonds to metal ions in dashed yellow, clashes are indicated in dashed red. The metal center is highlighted with asterisks. Interactions between ATP and the backbone of a residue are indicated with a yellow prime.

Using the same *in silico* approach ATP, phytate and PEP were docked into the active sites of the NtPAP, AtPAP12 and AtPAP26 models, respectively (Figs 6-8; **Table 2**). Phytate binds to NtPAP with a MolDock/Rerank score of -236.0/-132.2 kcal/mol. For the best poses of ATP and PEP, docking scores of -253.8/-170.2 and -169.6/-94.5, respectively, were obtained, consistent with the observations that NtPAP is able to effectively hydrolyze each of these substrates. Superposition of the rkbPAP crystal structure onto the NtPAP-phytate complex indicated that phytate is too bulky to be accommodated by rkbPAP in a similar pose. The comparison identifies His295 and Asp369 as the crucial residues that prevent phytate binding to rkbPAP. In NtPAP (and other phytases among the PAPs) these two residues are replaced by alanines (Ala374 and Ala437 in NtPAP), which enables a phosphate group from phytate to access a small, positively charged binding pocket in the enzyme (**Fig. 6**). In addition, the replacement of Tyr365, Gly366 and Val367 in rkbPAP by Arg433, Glu434 and Lys435, respectively, in NtPAP facilitates strong interactions with another two phosphate groups from phytate. Furthermore, residue Lys337 is conserved amongst the phytases aligned in this study (Arg258 in rkbPAP; **Figs 3** and **6**) and also interacts with a phosphate group from phytate (**Fig. 6**). The comparison between the rkbPAP structure and the NtPAP model thus indicates that all of the amino acid variations relevant to the absence (rkbPAP) and presence (NtPAP) of phytase activity are located in the C-terminal region of their polypeptide chains. Specifically, these residues define a conserved motif within the phytase group that is defined by the residues Arg433, Glu434, Lys435 and Ala437 (*i.e.* the “REKA” motif). It is noteworthy to add that Arg433 in NtPAP is predicted to interact with the carboxylate group of PEP in the NtPAP-PEP complex, thus presenting a likely structural element necessary to promote PEPase activity.

**Table 2.** Docking scores for the predicted complexes formed by rkbPAP, NtPAP, AtPAP12 and AtPAP26 with various substrates. Ligand efficiency scores are also shown.

| Substrate | Docking score (kcal/mol) | rkbPAP | NtPAP | AtPAP26 | AtPAP12 |
| --- | --- | --- | --- | --- | --- |
| **ADP** | Moldock* | -223.3 | ND | ND | ND |
|  | Rerank** | -150.0 |  |  |  |
|  | LE^ | -8.3 |  |  |  |
| **ATP** | Moldock | -337.7 | -253.8 | -189.0 | -211.4 |
|  | Rerank | -244.9 | -170.2 | -73.3 | -117.3 |
|  | LE | -10.9 | -8.2 | -6.1 | -6.8 |
| **Phytate** | Moldock | -192.6 | -236.0 | -79.8 | -200.9 |
|  | Rerank | +26.2 | -132.2 | +231.2 | -54.2 |
|  | LE | -5.4 | -9.8 | -2.2 | -5.6 |
| **PEP** | Moldock | -184.9 | -169.6 | -152.9 | -150.2 |
|  | Rerank | -122.4 | -94.5 | -113.4 | -91.2 |
|  | LE | -18.5 | -17.0 | -15.3 | -15.0 |

\*The binding energy of the ligand after post-processing of poses. \*\*The reranking algorithm uses a more advanced scoring scheme than the docking scoring function. Reranking will often improve the accuracy of the ranked poses in a prediction. ^MolDock Score divided by the heavy atom count.

**Figures 6:**
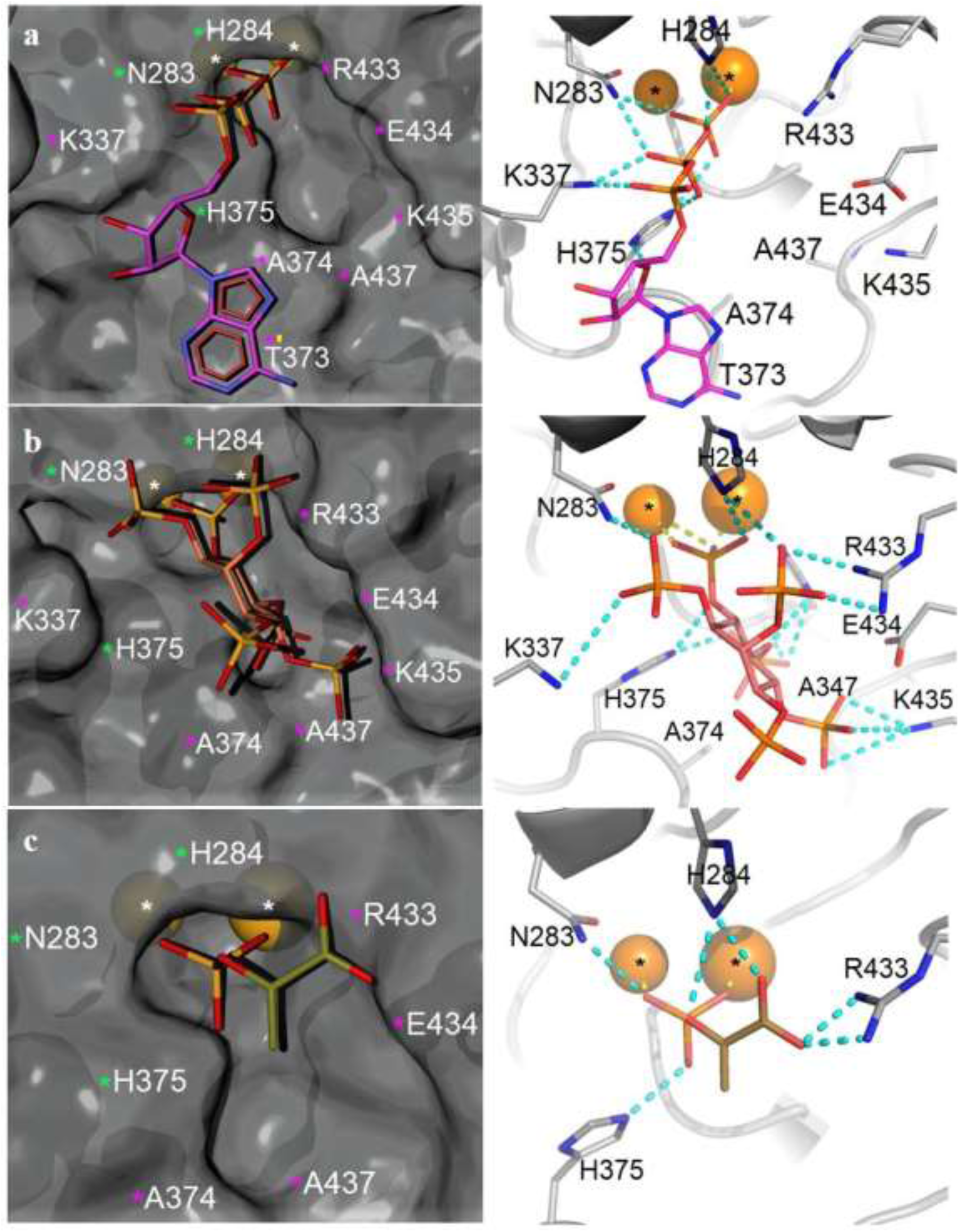
(left panel) Surface and (right panel) stick representations of the docking predictions of the binding modes of ATP (magenta carbons), phytate (salmon carbons) and PEP (brown carbons) to NtPAP (gray surface or carbons). Interactions between a substrate and the backbone of a residue are indicated with a yellow prime. Hydrogen bonds are indicated in dashed cyan and bonds to metal ions in dashed yellow. The metal center is highlighted with asterisks. Residues that have been mutated relative to the crystal structure of rkbPAP are marked with a magenta asterisk and residues that were not mutated are marked with a green asterisk. Interactions between ATP and the backbone of a residue are indicated with a yellow prime.

**Figures 7:**
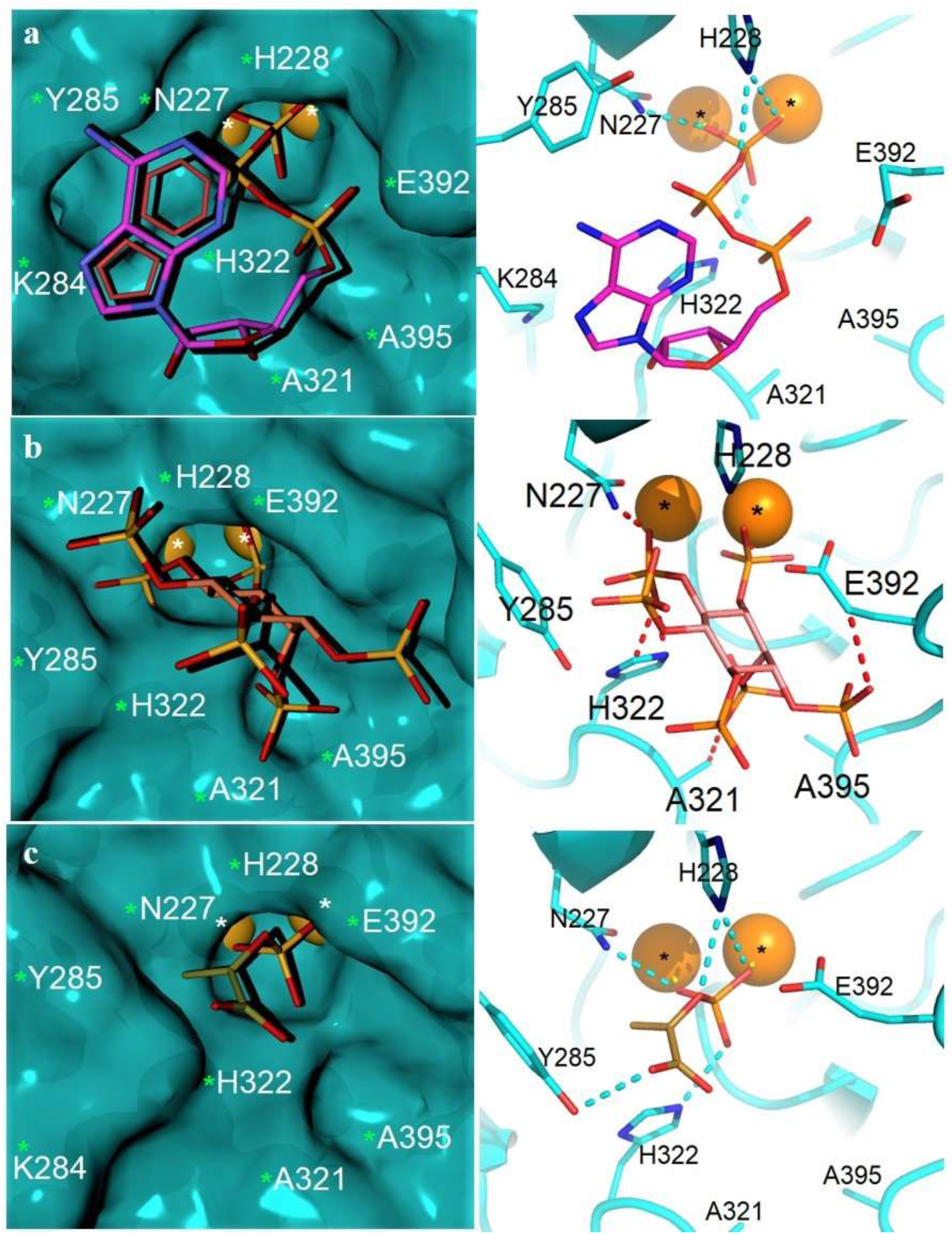
(left panel) Surface and (right panel) stick representations of the docking predictions of the binding modes of ATP (magenta carbons), phytate (salmon carbons) and PEP (brown carbons) to AtPAP26 (cyan surface or carbons). Hydrogen bonds are indicated in dashed cyan and bonds to metal ions in dashed yellow, clashes are indicated in dashed red. The metal center is highlighted with asterisks. Due to the high sequence similarity between AtPAP26 and IbPAP2 no residues in the active site had to be mutated in the modeling process. Conserved residues outside the immediate metal binding center are marked with a green asterisk.

**Figures 8:**
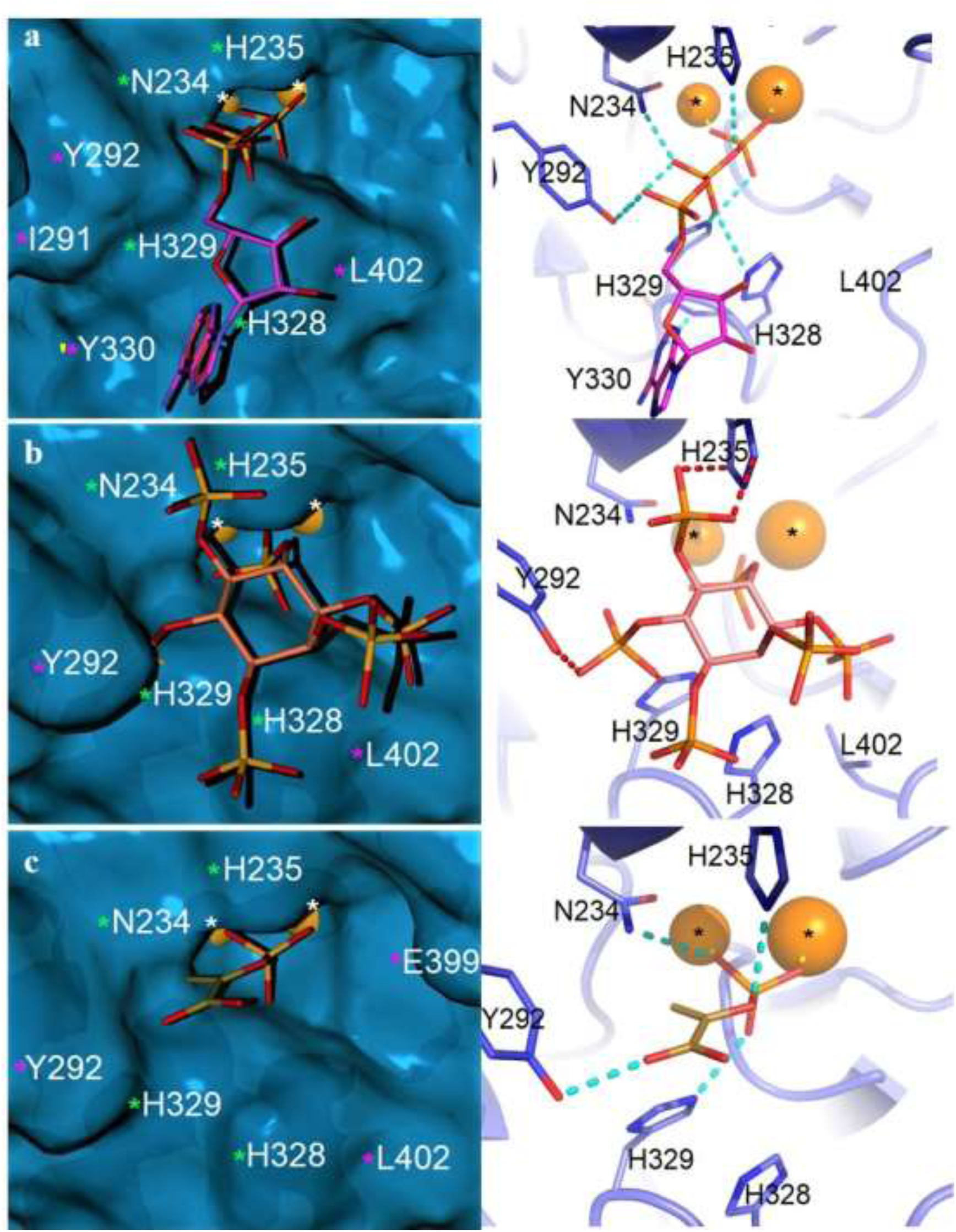
(left panel) Surface and (right panel) stick representations of the docking predictions of the binding modes of ATP (magenta carbons), phytate (salmon carbons) and PEP (brown carbons) to AtPAP12 (blue surface or carbons). Interactions between a substrate and the backbone of a residue are indicated with a yellow prime. Hydrogen bonds are indicated in dashed cyan and bonds to metal ions in dashed yellow, clashes are indicated in dashed red. The metal center is highlighted with asterisks. Residues that have been mutated relative to the crystal structure of rkbPAP are marked with a magenta asterisk and residues that were not mutated are marked with a green asterisk. Interactions between ATP and the backbone of a residue are indicated with a yellow prime.

For the AtPAP26 model the docking scores for the optimal poses of phytate and PEP were -79.8/231.2 kcal/mol and -152.9/-113.4 kcal/mol, respectively, with phytate having a very poor Rerank score, consistent with various clashes within the active site (**Fig. 7**). In the AtPAP26-PEP complex the phosphate group of the substrate binds directly to the two metal ions in the active site and is further stabilized via hydrogen bonds to residues His322 and His228. The alkene group of PEP slots into a pocket formed between the aromatic group of Tyr285 and the sidechain of Asn227, while the hydroxyl group of Tyr285 is predicted to form a hydrogen bond with the carboxylate group of PEP (**Fig. 7**). Superposition of the AtPAP26-PEP and NtPAP-phytate complexes indicates that in particular the close proximity of the side chains of Tyr285 and Glu392 to the metal ion center in AtPAP26 results in a sequestered active site that easily accommodates PEP but prevents phytate from binding. We thus predict that AtPAP26 is not capable of hydrolyzing phytate. This enzyme is also reported to be a poor ATPase (see above). It was thus not surprising that the majority of obtained poses with ATP had positive Rerank scores and many clashes within the active site were observed. However, in one pose the docking score was favorable (-189.0/-73.3 kcal/mol). In this pose the terminal phosphate group of ATP binds bidentately to the metal center, but both Tyr285 and Glu392 need to move significantly to accommodate the remaining two phosphate groups of the substrate (**Fig. 7**). This pose may thus reflect the ability of AtPAP26 to hydrolyze ATP, albeit poorly [72].

In contrast to AtPAP26, AtPAP12 is a PEPase with good ATPase activity [12], in agreement with its phylogenetic proximity to other plant PAPs that are efficient ATPases (*i.e.* AtPAP10 and rkbPAP [43, 61]; **Fig. 3**). Phytate, ATP and PEP were docked into the active site of AtPAP12, and the relevant docking scores are -200.9/-54.2 kcal/mol, -211.4/-117.3 kcal/mol and -150.2/-91.2 kcal/mol, respectively. For phytate, the large difference between the MolDock and Rerank scores again suggests unfavorable binding interactions, and indeed various clashes with several active site residues are evident (**Fig. 8**). It is thus unlikely that AtPAP12 is able to hydrolyze phytate. The docking scores for PEP are similar to those of AtPAP26, in agreement with the similar PEPase activities recorded for these two enzymes. Furthermore, for ATP the docking scores are comparable to those obtained for the NtPAP-ATP complex, consistent with the proficient ATPase activity of AtPAP12. His295/His328 in rkbPAP/AtPAP12 forms a hydrogen bond with an oxygen atom of one of the phosphate groups of the substrate (**Fig. 8**). However, in AtPAP26 the corresponding residue is an alanine (Ala321), which may thus weaken the interaction with ATP.

The observed sequence variations that link specific plant PAP variants to characteristic substrate preferences are useful in assigning functions to novel members of this group of enzymes. The residues identified as “markers” for particular substrate specificities are summarized in **Table 3**, and are also exemplified in a comparison of three PAP sequences identified in *Eucalyptus grandis* (*i.e.* EgPAP2X1, EgPAP26 and EgPAP15; Figs. 3 and **S1**). Although the eucalypt PAPs have not been characterized, it is evident that, similar to *A. thaliana*, rice and others, multiple forms of PAP are present. Based on their varying substrate specificities, the eucalypt PAPs may be involved in a diverse range of metabolic functions. Considering the oligotrophic environments most eucalypt species have adapted to (including many with very low phosphorus availability), PAPs from these plants and other wild plants adapted to low phosphorus environments may form ideal starting points for the engineering of PAP variants with desired substrate preference and stability to be used in applications related to increasing the phosphorus use of crops.

**Table 3.** Amino acid residues that influence the substrate preference of plant PAPs (highlighted in green). The residue numbers are according to the rkbPAP sequence. Also shown are residues that play an important role in substrate binding irrespective of the identity of the reactant (in purple). In particular with respect to phytate binding a motif in the C-terminal region (“REKA”) is important (residues in bold).

| Residue position | 201 | 202 | 258 | 259 | 295 | 296 | 365 | 366 | 367 | 369 |
| --- | --- | --- | --- | --- | --- | --- | --- | --- | --- | --- |
| rkbPAP | H | H | R | G | H | N | Y | G | V | D |
| AtPAP12 | H | H | I | Y | H | N | S | E | G | L |
| AtPAP26 | H | H | K | Y | A | N | Q | E | G | A |
| NtPAP | H | H | K | S | A | N | **R** | **E** | **K** | **A** |
| GmPHY | H | H | K | T | A | N | **R** | **E** | **K** | **A** |
| OsPHY1 | H | H | K | S | A | N | **R** | **E** | **K** | **A** |

## Conclusions

In conclusion, in this study a simple assay was developed that facilitates the assessment of the substrate specificity of PAP, a metalloenzyme that has numerous copies in many plants. Using this assay, it was demonstrated that rkbPAP is both a potent ATP- and ADPase, but it has no activity against phytate. The crystal structure of rkbPAP in complex with a vanadate derivative of ATP visualizes relevant binding interactions between this substrate and the enzyme. Based on available crystal structures of plant PAPs models of NtPAP, AtPAP12 and AtPAP26 were generated, PAPs that differ in their substrate preferences. Docking studies with phytate, PEP and ATP as substrates resulted in the identification of a number of active site residues that play a crucial role in determining substrate preference. In the context of applying PAP in agricultural biotechnology, the discovery of the REKA motif in PAPs with phytase activity (NtPAP, OsPHY1 and GmPHY) is of particular relevance as it facilitates the engineering of PAP variants with a preference for the important nutrient phytate while maintaining strong nucleotidase and PEPase activity. PAPs identified in the fast growing *E. grandis* and other efficient P using species may play an important role in such endeavors.

## Acknowledgements

The authors acknowledge funding from the Australian Research Council (DP0986292). Preliminary X-ray data were measured at the University of Queensland Remote Operation Crystallisation and X-ray diffraction facility (UQROCX). The final X-ray data were obtained at the Australian Synchrotron. The assistance of Tom Caradoc-Davies and Alan Riboldi-Tunnicliffe during data collection was much appreciated.

## Author contributions

DF purified and crystallized the enzyme, and together with LWG collected the X-ray data and refined crystal structures. DF also interpreted structural data and performed *in silico* simulations. TL performed the ITC experiments in the supplementary section. The crystal soaking experiment was designed and performed by DF. All authors contributed to the writing of the manuscript, and LWG, GS and RPM were responsible for the experimental design and overall supervision of the project progress and manuscript writing.

## Footnotes

1 We cannot unambiguously rule out that the bound ligand may be a mixture of species (*i.e.* ADPV, AMP-divanadate (AMPDV) and adenosine-trivanadate (ATV)). ADPV, AMPV and AMPDV are known to form spontaneously in solution from ADP, AMP and vanadate [68]. However, the bond length refinements and resulting electron maps in the structure presented here fit ADPV better than other species. Furthermore, irrespective of the precise nature of the bound ligand it provides unprecedented insight into the binding mode of a complex endogenous substrate-like molecule to a PAP.

