## Supplemental data for "Structural elements that modulate the substrate specificity of plant purple acid phosphatases: avenues for improved phosphorus acquisition in crops"

*Experimental details for the ITC assay*. The enthalpy change ($\Delta H$) associated with the reaction of rkbPAP with *p*-NPP was determined by injecting 10 µL of *p*-NPP (20 mM) into the reaction cell (200 µL) containing 0.95 µM of rkbPAP, and by allowing the reaction to proceed to completion. The reaction is exothermic as indicated by the negative value of the heat pulse (dQ/dT); the value of $\Delta H$ can be calculated by integrating the area under the curve in **Fig. S2a** (minus the endothermic heat associated with the dilution) divided by the amount of *p*-NPP hydrolyzed in the cell. The value for the endothermic heat generated by the dilution was obtained by injecting 10 µL of *p*-NPP (20 mM) into the reaction cell containing only buffer: $\Delta H$ = 3089 ± 52 µcal/µmol (*i.e.* 12.9 ± 0.22 kJ/mol), in good agreement with the corresponding $\Delta H$ value determined by enzymatic assays (13.2 ± 0.18 kJ/mol) [1]. In this study, we focused on measuring initial catalytic rates in order to minimize the effect of the reaction product and competitive inhibitor phosphate [2–6]. Different concentrations of *p*-NPP were prepared and 8 µL of 0.24 µM rkbPAP were injected into the substrate solutions. **Fig. S2b** shows a typical calorimetric trace of a reaction. The initial rates determined in µcal/s, when divided by $\Delta H$ (µcal/µmol), are transformed into µmol/s, representing the amount of *p*-NPP hydrolyzed per second in the cell. These rates were subsequently plotted against the *p*-NPP concentration and fitted to the Michaelis-Menten equation (Eq. 1; **Fig. S2c**), resulting in a $k_{cat}$ value (*i.e.* $V_{max}$/[PAP]) of ~190 $s^{-1}$ and a Michaelis constant $K_m$ of ~4 mM.

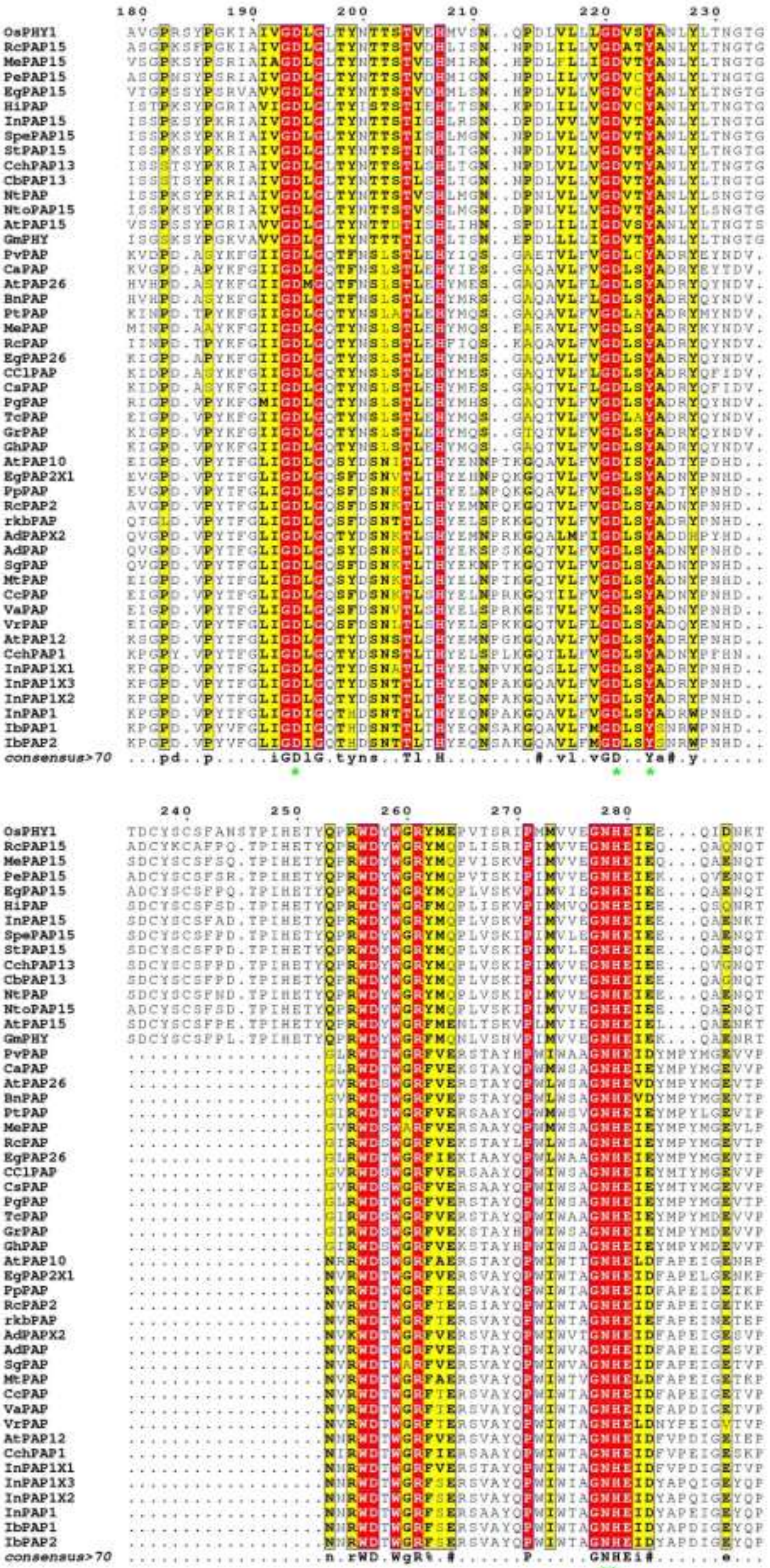

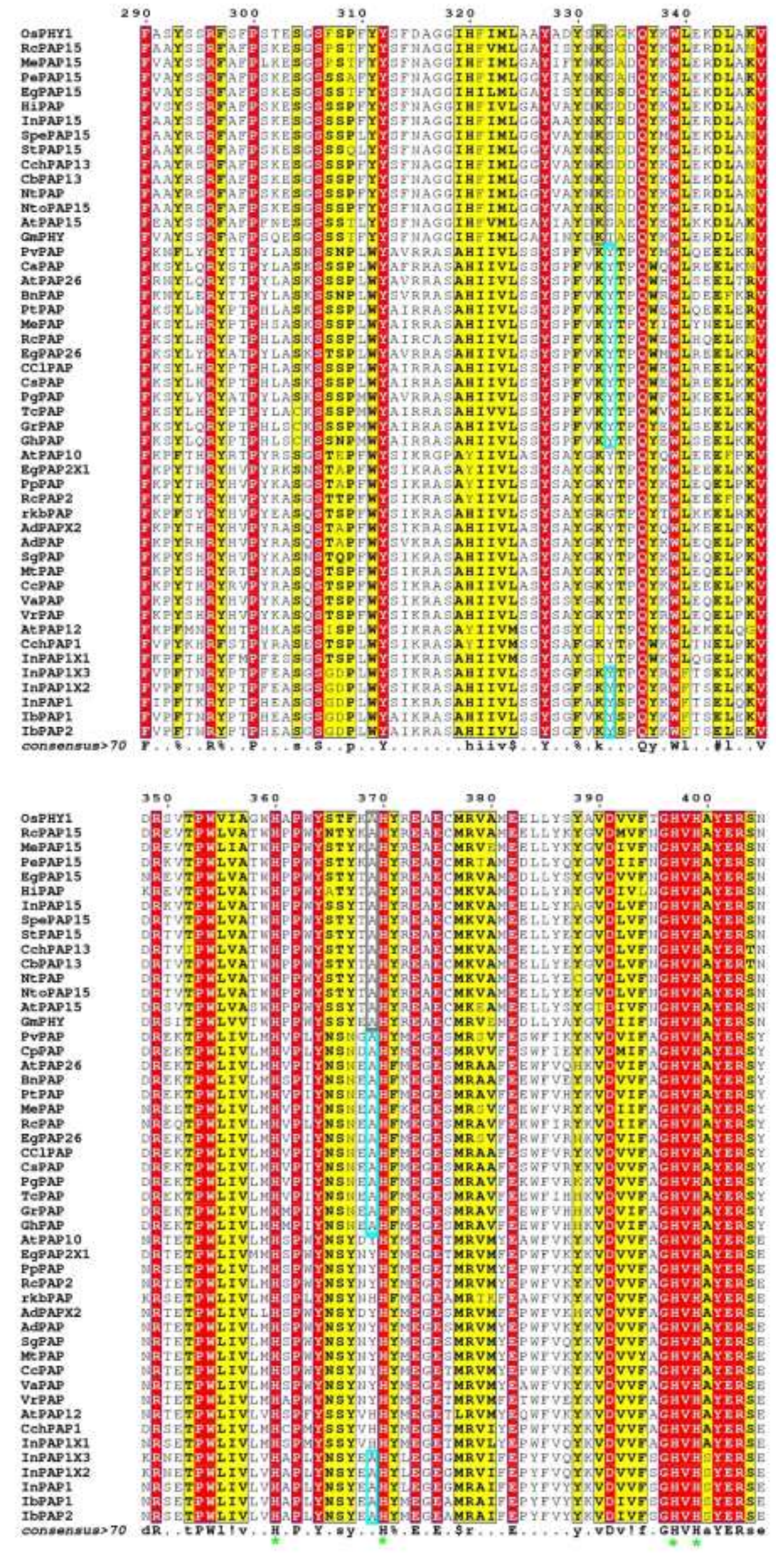
290 300 310 320 330 340
OsPHY1
RcPAP15
MePAP15
PePAP15
EgPAP15
HiPAP
InPAP15
SpePAP15
StPAP15
CchPAP13
CbPAP13
NtPAP
NtoPAP15
AtPAP15
GmPHY
PvPAP
CaPAP
AtPAP26
BnPAP
PtPAP
MePAP
RcPAP
EgPAP26
CClPAP
CsPAP
PgPAP
TcPAP
GrPAP
GhPAP
AtPAP10
EgPAP2X1
PpPAP
RcPAP2
rkbPAP
AdPAPX2
AdPAP
SgPAP
MtPAP
CcPAP
VaPAP
VrPAP
AtPAP12
CchPAP1
InPAP1X1
InPAP1X3
InPAP1X2
InPAP1
IbPAP1
IbPAP2
consensus>70 F..%..R%..P....s.S..p..Y.......hiiv$..Y..%.k...Qy.Wl..#l..V
350 360 370 380 390 400
consensus>70 dR..tPWl!v..H.P.Y.sy..H%.E.E.$r...E.....y.vDv!f.GHVHaYERse

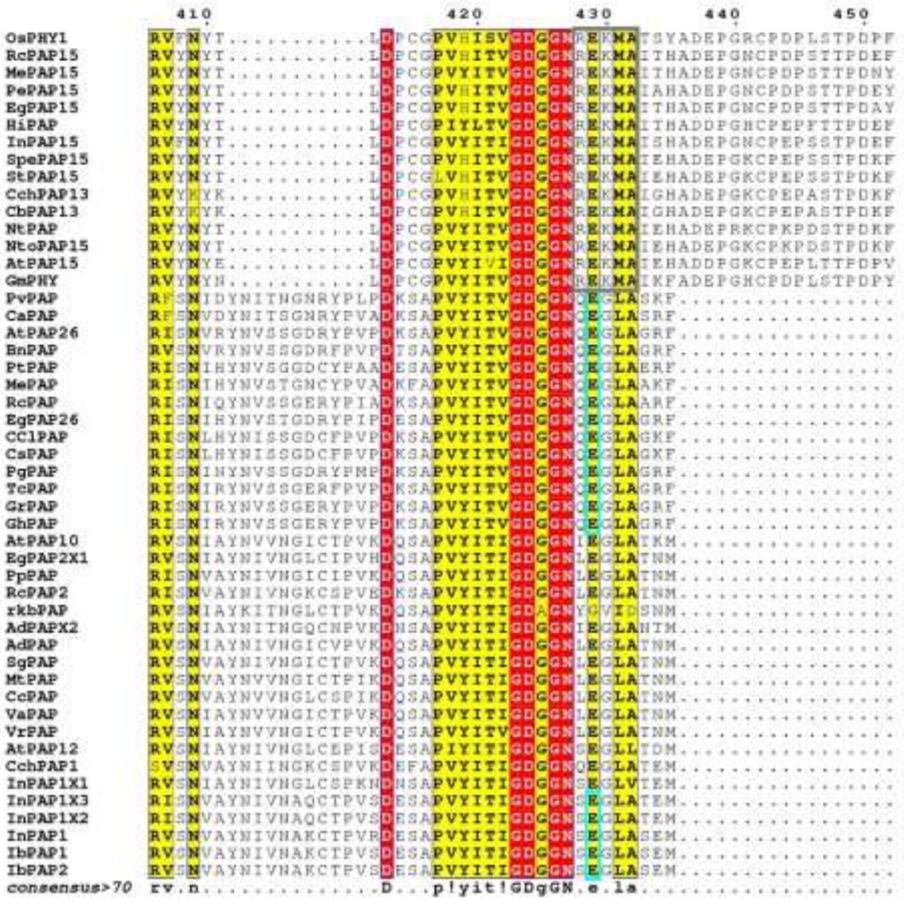

**Figure S1**: Clustol-O Multiple sequence alignment of hypothetical and known plant PAPs, displayed using ESPript 3.0. Fourty nine PAPs from agriculturally relevant crops were aligned. Conserved elements are in red and semi-conserved elements in yellow. Consensus numbering and sequence are shown above and below the alignment, respectively. Metal-coordinating residues and the two histidine residues that form the core active site of PAPs (**Fig. 1**) are marked with a green asterisk. Conserved residues characteristic of PEPases and phytases are boxed in cyan and gray, respectively.

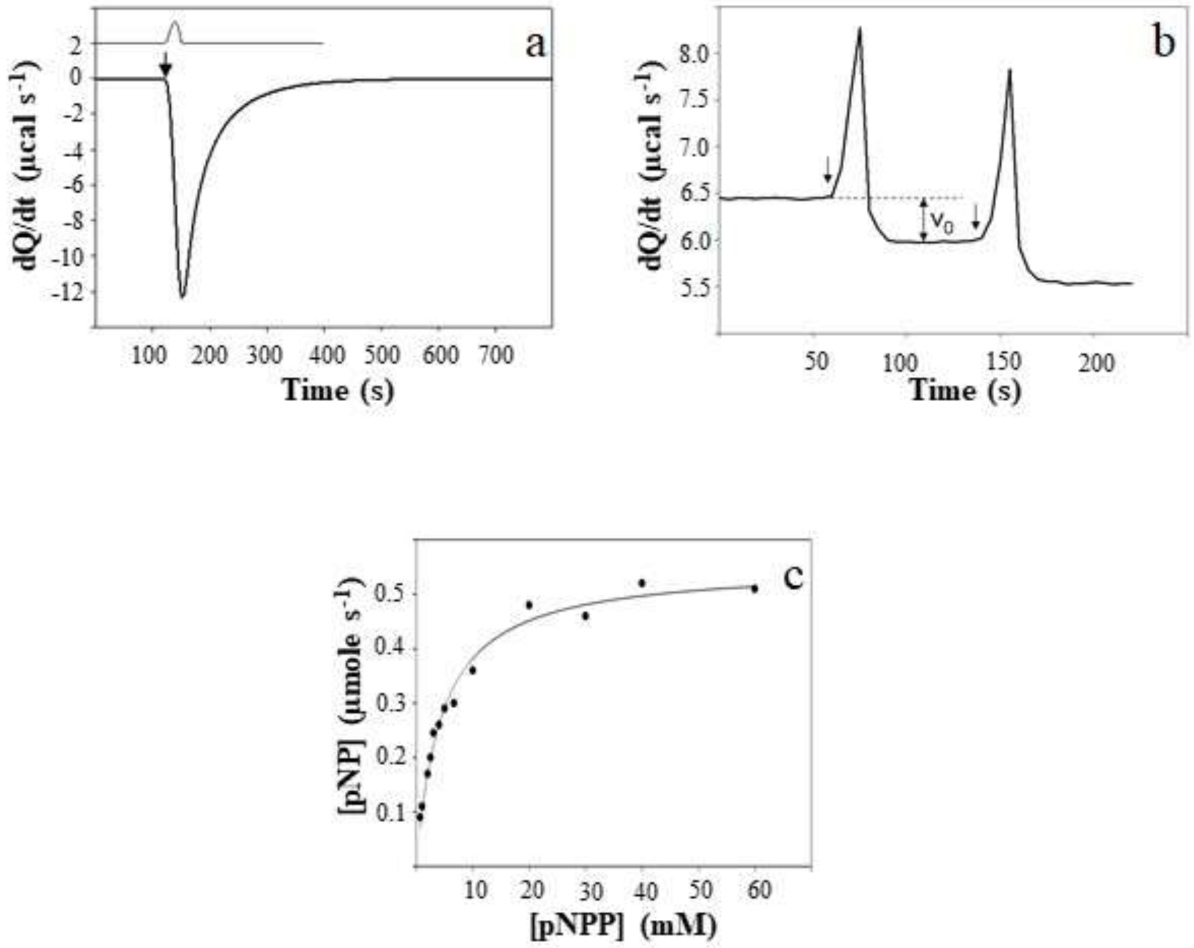


**Figure S2:** Establishment of an ITC-based catalytic assay to measure the enzymatic activity of PAPs. (a) The hydrolysis of *p*-NPP by rkbPAP is exothermic. (b) A typical data set of two subsequent injections of *p*-NPP into the rkbPAP solution. (c) The experimental data were converted into the amount of *p*-NPP hydrolyzed per second, and fitted to the Michaelis-Menten equation to calculate the values of $k_{cat}$ and $K_m$ of this reaction. Details of the data analysis are described elsewhere [7].

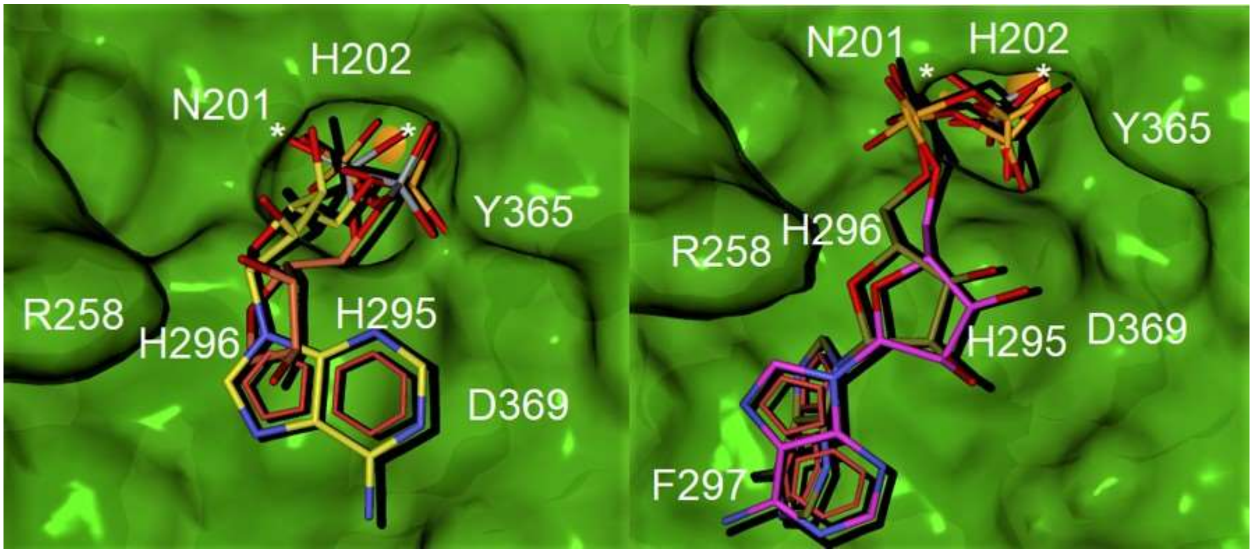


**Figure S3**: A comparison of docking predictions and crystal structures. Docking prediction of the binding modes of (left) ADP (yellow carbons) and (right) ATP (brown carbons) to rkbPAP (green surface). The corresponding crystal structures of rkbPAP in complex with ADV (salmon carbons) (6HWR [8]) and ADPV (magenta carbons) are superimposed on the docking predictions. Metal ions are shown as orange spheres and marked with an asterisk.
